# Nanopore event detection in a simple and adaptive way

**DOI:** 10.64898/2026.05.07.723187

**Authors:** Peijia Wei, Mayukh Kansari, Michael Mierzejewski, Tobias Ensslen, Chih-Yuan Lin, Kyril Kavetsky, Peter D. Jones, Jan C. Behrends, Marija Drndić, Maria Fyta

**Affiliations:** Computational Biotechnology, RWTH Aachen University, Worringerweg 3, 52074 Aachen, Germany; NMI Natural and Medical Sciences Institute at the University of Tübingen, 72770 Reutlingen, Germany; Laboratory for Membrane Physiology and Technology, Department of Physiology II, Faculty of Medicine, University of Freiburg, Hermann-Herder-Str. 7, 79104 Freiburg, Germany; Hahn-Schickard Institute for Microanalysis Systems, Georges-Koehler Allee 304, 79110 Freiburg i.Br.; Department of Physics and Astronomy, University of Pennsylvania, Philadelphia, Pennsylvania 19104, United States; Center for Computational Life Sciences, RWTH Aachen University, Pauwelstrasse 19, 52074 Aachen, Germany

## Abstract

Nanopore read-out, that is the current signals measured across nanometer-sized openings in dielectric membranes or through natural protein channels, enables the detection, identification and sequencing of individual molecules. The detection can take place by analyzing the events of single biomolecules interacting with the pore. The accuracy in the detection of these single events is key for identification of physicochemical properties of analyte molecules. To this end, we further develop a very simple, fast, almost parameter-free, and adaptable cluster-based event detection (CBED) algorithm that clusters the nanopore signals prior to detecting nanopore events. The algorithm is validated against two other event detection schemes with respect to simplicity and efficiency. For this, nanopore data from four different experiments stemming from different laboratories that vary in the nanopore type, size, and analyte are considered. The comparison is made on the basis of the number of events detected, their quality, and the most important features extracted from nanopore events. Our results underline the higher efficiency and less noise of the CBED detected events for biological nanopore data and the need for an on-the-fly adaptivity of the baseline current for a class of solid-state nanopore data.

## I. INTRODUCTION

Nanopores are orifices of molecular size in either dielectric material or channel proteins, through which analyte molecules can be driven by electrodiffusion, electroosmosis, and local electrophoretic forces [1–5]. This passage can be monitored through modifications in the ionic (longitudinal) [6, 7] and/or the electronic (transverse) current [8, 9] across the nanopore. As molecules pass through or interact with the pore, even without full translocation [10, 11], they partially block the current, producing characteristic signatures in the measured signal, known as current blockades. These distinct events differ from the baseline current of an open nanopore guiding only ions of the solution. The nanopore current measurements, including baseline and blockade signals, are time traces characterized by distinct events reflecting molecular translocations, sequence variations, or interactions with the nanopore [12–14]. These current signals provide rich information about the physical and chemical properties of the analyte, including its size, charge, conformation, or sequence [15]. Consequently, nanopore sensing has emerged as a powerful platform for single-molecule detection and sequencing applications [16].

In order to read-out and interpret the raw ionic current signals, various computational approaches and signal processing algorithms have been proposed, targeting enhanced accuracy, sensitivity, and robustness of molecular detection [17–22]. Read-out algorithms for nanopore sequencing data typically involve several crucial stages, including signal preprocessing, noise reduction, feature extraction, event detection, and subsequent event classification or identification [23]. Despite significant progress in nanopore signal processing, it is still difficult to accurately detect individual events in nanopore signals. This challenge stems from the complexity of the ionic current, which depends on the shape and functionalization of the nanopore, molecular dynamics, background noise, noise interference, baseline drift, and overlapping event signatures [24–26].

Event detection methods like the cumulative sum (CUSUM) [27] and the Pruned Exact Linear Time (PELT) [28] algorithm based on different schemes and parameterization are commonly used. CUSUM is a sequential, real-time method highly dependent on tuning parameters such as the decision threshold, requiring user expertise for optimal settings [29]. Similarly, PELT is an offline approach, sensitive to the appropriate choice of penalty parameter, and often requires rigorous empirical adjustments [28]. Beyond these, adaptive thresholding has become a standard strategy to compensate for baseline drift and noise in nanopore signals. Several end-to-end nanopore analysis schemes, including OpenNanopore [29], Nanopore Analysis [30], Transalyzer [31], MOSAIC [32], EventPro [33], and PyPore[34], employ per-sample real-time estimates of the baseline mean and standard deviation for event detection. Most recently, the Dynamic Correction Method [35] has further improved baseline tracking and threshold adjustment via per-sample low-pass filtering of local statistics. Additionally, Bayesian frameworks employing Gaussian mixture models (GMM) have demonstrated robust detection performance, especially in handling diverse and noisy signal conditions. These probabilistic models can infer underlying signal states with minimal user intervention but typically require careful tuning of prior assumptions [36]. Building on this, hidden Markov model (HMM)-based detection algorithms have also been proposed, where temporal dynamics are incorporated through state transitions, enabling more accurate decoding of signal segments using methods such as the Viterbi algorithm [37–39]. These models are independent of explicit thresholding by inferring event sequences probabilistically and have shown improved performance over GMM-based approaches, particularly in noisy environments [40].

In this study, we analyze data from four distinct nanopore experiments and compare different event detection algorithms. For this, we propose the use of the cluster-based event detection (CBED) method and its adaptive version and systematically evaluate this together with two of the most common event detection schemes (CUSUM and PELT). Our enhanced implementation of CBED leverages a histogram-based, two-component Gaussian mixture model to identify and distinguish baseline and event clusters, from which an adaptive detection threshold is directly derived. Recognizing scenarios where baseline signals drift significantly or become indistinct from events, we extend CBED to include an adaptivity module, which employs sliding-window adjustments to both baseline estimation and thresholding, akin to the recent adaptive thresholding strategy [35]. Unlike traditional algorithms that require intensive parameter tuning, CBED, particularly in its adaptive form, significantly reduces the necessity for empirical adjustments.

Through comprehensive analysis on diverse nanopore datasets, including biological nanopores (with events, clearly distinguishable from the baseline) and solid-state nanopores (with low signal-to-niose ratio and drifting baselines), we demonstrate the efficiency, reduced noise sensitivity, adaptability, and ease-of-use of our CBED algorithm. Our contribution specifically lies in refining existing cluster-based methods through practical adaptations, optimizing event-calling accuracy under varying experimental conditions, and minimizing user-defined parameter dependency, thereby providing a robust and user-friendly computational tool for nanopore signal analysis.

### II. METHODOLOGY

### A. Review of event detection algorithms

In this study, we aim to both explore and extend our CBED event detection algorithm also in comparison to the CUSUM and Pelt schemes mentioned above. In order to facilitate an in depth comparison instead of pure visualization, we briefly discuss the main mathematical details of these two algorithms. Rough event localization is performed as an essential preprocessing step before applying event detection algorithms. This step aims to identify regions within the ionic current signal that are likely to correspond to an event. By restricting the analysis to these localized regions, the computational cost of the subsequent process is significantly reduced, while minimizing the risk of falsely identifying signal fluctuations within the baseline noise as true events. A common approach for rough event localization is based on a global threshold, where events are detected if the current drop exceeds *σ*_thresh_, a multiple of the baseline noise level:

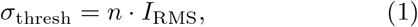

with *I*_RMS_ being the standard deviation of the baseline current and *n* a user-defined sensitivity parameter.

For signals exhibiting low-frequency baseline fluctuations, a local thresholding method is applied. Here, the baseline mean and standard deviation are continuously estimated over a moving window of previous data points, and dynamic thresholds for event start *η*_*S*_(*k*) and end *η*_*E*_(*k*), respectively are computed as:

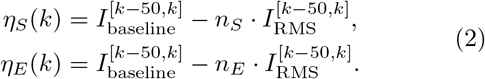

Where 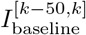 and 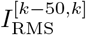denote the local mean and standard deviation of the baseline calculated over the previous 50 data points. After rough event localization, the extracted signal segments can be subjected to subsequent event detection using the CUSUM or PELT algorithms to identify abrupt changes and segment the events into discrete levels.

The CUSUM algorithm, first introduced in 1954, is a sequential analysis technique widely used for change-point detection in time series [41]. This method is categorized as an online approach, wherein change points are detected in real time as data points are sequentially processed, rather than by analyzing the complete signal afterwards. The CUSUM algorithm detects abrupt changes in the mean of nanopore ionic current signals. For a signal segment starting at index *k*_0_, the mean *m*_*k*_ and variance *v*_*k*_ are iteratively computed as:

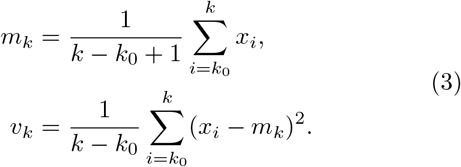

with *k* the segment considered and *x*_*i*_ the current value at index *i*. The instantaneous log-likelihood for current sample assuming local baseline has moved in the positive direction (*s*_*p*_[*k*]) or negative direction (*s*_*n*_[*k*]) are calculated for each sample *x*[*k*]:

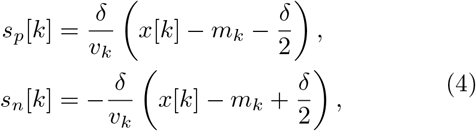

where *δ* is the expected mean shift magnitude. Cumulative sums *S*_*p*_[*k*] and *S*_*n*_[*k*] are updated as:

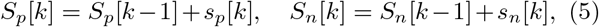

with the accumulate positive and negative decision function defined respectively as:

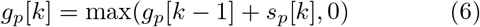

and

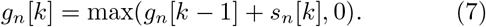

A change from baseline to blockade or vice versa is detected if *g*_*p*_[*k*] *> h* or *g*_*n*_[*k*] *> h*, where *h* is a predefined threshold. The exact transition is identified by minimizing *S*_*p*_ or *S*_*n*_ within the segment. Parameters *δ* and *h* are served as the decision threshold for the log-likelihood ratio.

The PELT algorithm is an offline method that segments signals into statistically homogeneous intervals by identifying abrupt changes [28]. The algorithm operates by minimizing a cost function *V* (*T*), defined as the sum of individual segment cost functions *c*(*y*_*a,b*_), reflecting the homogeneity within each segment delimited by change points. For signals assumed to follow a Gaussian distribution, this cost function takes the form:

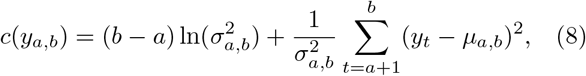

where 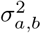 and *µ*_*a,b*_ represent the variance and mean of segment *y*_*a,b*_, respectively with *a* and *b* the start and end index of the segment. The sensitivity of detecting a change from baseline to blockade current or vice versa is controlled by incorporating a penalty term pen(*T*), typically expressed as:

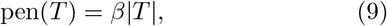

where |*T*| denotes the number of detected change points (from baseline to event and vice versa) and *β* is a parameter that adjusts the detection sensitivity. Higher *β* values require stronger evidence for accepting a change point, reducing false detections. For an automated, parameter-free selection of *β*, the Bayesian Information Criterion (BIC) [42] can be applied to achieve an optimal balance between model complexity and data fit quality, i.e. quality of the empiricity in the model.

### B. Cluster-based event detection (CBED)

Having addressed the main concepts of CUSUM and Pelt, we introduce the basic details of the cluster-based event detection (CBED) algorithm. CBED proceeds in four stages: (1) Clustering the current sample on the basis of cluster distributions, (2) threshold determination (3) event calling and (4) post-processing. A variant using a fixed global baseline is described first, followed by an extension for variable baselines rendering CBED adaptive to baseline shifts. In order to clarify the sequence of operations, we provide a workflow diagram (Fig. 1).

**FIG. 1:**
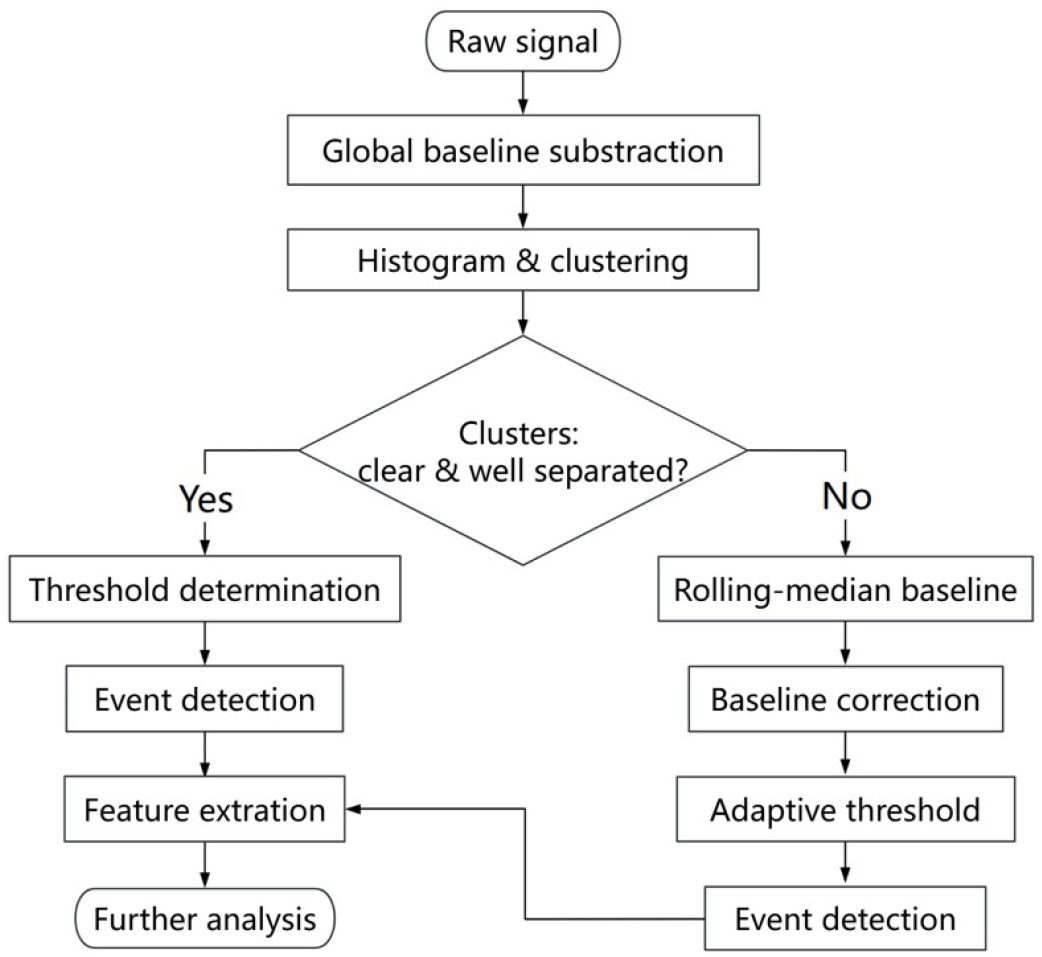
The adaptable CBED event detection algorithm: the workflow taking the raw nanopore data and performing the analysis up to the event detection. The labels are self-explaining (see text for details).

#### Histograms and Clustering

We begin by constructing the empirical histogram of the baseline-corrected current samples 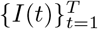. We then approximate its probability density by a two-component Gaussian mixture model:

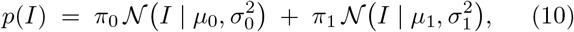

with 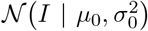 the Gaussian distributions, *p*(*I*) the current distributions and the mixture weights satisfy *π*_0_, *π*_1_ ≥0 and *π*_0_ + *π*_1_ = 1. Each component density is given by

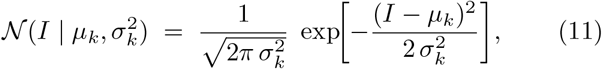

with *k*∈{0, 1}. The component with weight *π*_0_ (mean *µ*_0_, variance 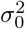) is identified as the open pore (baseline) cluster, while the smaller-weight component 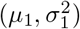 represents blockade events.

In the clustering step, we approximate the empirical distribution of current samples by a convex combination of two Gaussian densities. The overall model is a “mixture” of two Gaussians, one representing the open pore (baseline) cluster (*µ*_0_, *σ*_0_) and the other the blockade (event) cluster (*µ*_1_, *σ*_1_), hence it is is a *Gaussian mixture model*. This formulation naturally captures both the overlap between baseline and event currents and their relative prevalence via the weights *π*_0_ and *π*_1_. In practice, we approximate this model using a two-component K-Means clustering algorithm [43], where the resulting cluster centroids *µ*_0_ and *µ*_1_ serve as the means of the two Gaussian components. The empirical histogram of the current samples is then inspected to assess whether the two clusters are well separated or overlapping, which determines which detection strategy is applied subsequently.

#### Threshold Determination

The current signals considered as events are defined by simultaneously enforcing a *vertical* (amplitude) threshold and a *horizontal* (duration) threshold, as follows:

#### Vertical threshold

With the mixture parameters {*π*_*k*_, *µ*_*k*_, *σ*_*k*_*}* in hand, the detection threshold *ϵ*_*V*_ can be defined as the solution to the equality of the two component densities:

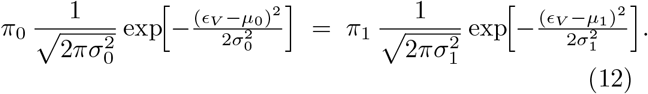

Taking the natural logarithm of both sides yields:

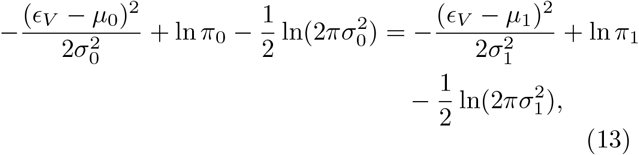

which can be rearranged into a quadratic equation in *ϵ*_*V*_. The root that lies between *µ*_0_ and *µ*_1_ corresponds to the “valley” of the mixture distribution and thus the optimal boundary between baseline noise and blockade events.

#### Horizontal threshold

Given a sampling rate *f*_*s*_ (Hz), a minimum event duration *τ*_min_ (s) translates to

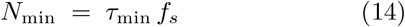

consecutive samples. We group contiguous indices in ℐ into *n* clusters {*t*_1_, …, *t*_*n*_} and retain only those for which

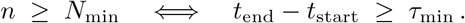

#### Event definition

An *event* is thus any contiguous interval [*t*_start_, *t*_end_] satisfying necessarily both

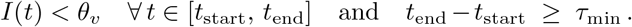

#### Adaptive Baseline and Threshold Adjustment

In the cases - typically observed in solid-state nanopore recordings - revealing gradual open-pore baseline drifts, leading to overlapping “baseline” and “event” clusters, we replace the clustering method with a sliding-window clustering procedure following the four steps:

1. Rolling-median baseline: Let the raw current trace be *I*(*t*) sampled at rate *f*_*s*_. Over a window of duration *W* (in seconds), corresponding to *N* = *W* ×*f*_*s*_ samples, we compute the rolling-median baseline

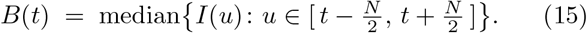 This nonparametric estimate is centered and uses at least one sample when the full window is unavailable.
2. Baseline correction: We subtract the local baseline from the raw trace,

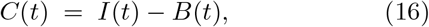

yielding a zero-centered residual *C*(*t*) whose distribution reflects only the fast fluctuations and blockade events.
3. Adaptive threshold: From the entire corrected trace {*C*(*t*)}, we compute its global median 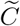 and standard deviation *σ*_*C*_. We then set the detection threshold

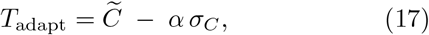

with *α* chosen empirically (here *α* = 1.5). By using the current residuals’ own statistics, the threshold self-adjusts to changes in noise level and slow baseline wander.
4. Event detection: We flag all timepoints *t* for which *C*(*t*) *< T*_adapt_ as candidate blockade samples. These indices are grouped into contiguous clusters by joining any two points separated by at most *δ*_max_ samples to form preliminary event intervals. Finally, we enforce a minimum event duration Δ*t*_min_ by discarding any cluster containing fewer than *f*_*s*_ × Δ*t*_min_ samples. This sliding-window, rolling-median scheme continuously adjusts both the baseline estimate and the detection threshold, ensuring high sensitivity to true blockade events while rejecting spurious fluctuations even in the presence of slow drift.

### C. Nanopore data

In order to compare the event detection schemes, CUSUM, PELT and our adaptable CBED algorithm, we use diverse types of nanopores data. This diversity refers to both the nanopore material and size, as well as the analyte type. Biological and solid-state nanopores often exhibit different blockade amplitudes due to differences in pore geometry and surface interactions [44–46]. To this end, we analyze the recordings from the following experiments for which the most relevant information is provided in Table I:

- homo-ssDNA either 5A-long or a mixture of 3A and 5A homo-DNA through a biological aerolysin pore. The buffer solution consists of 4 M KCl and 10 mM Tris at pH 7.5 [47]. The current signal was recorded at a sampling rate of 1 MHz with a final bandwidth of 33.3 kHz. We label these data as “A5” and “A3A5”, respectively.
- Single-stranded DNA homopolymer (Integrated DNA Technologies, Inc.) translocates through a 1.2 nm diameter nanopore (pore B in Ref. [48]). Data were recorded at a 10 MHz bandwidth. The buffer solution consists of 3 M KCl, 10 mM TrisHCl, and 1 mM EDTA at pH 8. We label these data as “ssDNA”.
- Peptide-oligonucleotide-conjugate (Biomers.net GmbH) translocates through a 3 nm diameter silicon-nitride nanopore. Data were recorded at a 10 MHz bandwidth (Elements srl) [49]. The buffer solution consists of 3 M KCl, 10 mM HEPES, and 10 mM MgCl_2_ at pH 7.4. We label these data as “POC”.
- *λ*-phage DNA (5 µg/mL) translocations through a solid-state silicon nitride nanopore (14 nm diameter, 50 nm thickness) using a microfluidic chip setup. The electrolyte conditions were asymmetric, with 1 M KCl on the *cis* side and 3 M KCl on the *trans* side. A holding voltage of 140 mV was applied, and the current signal was recorded at a sampling rate of 400 kHz with a 100 kHz low-pass filter. We label these data as “*λ*-DNA”.

The selected datasets span a range of signal-to-baseline contrasts (which is partly a function of the molecule sizes and amplifier bandwidth used): high contrast in A3A5 (biological nanopore), moderate contrast in *λ*-DNA, and low contrast in ssDNA and POC (last three samples in solid-state pores). High-contrast traces feature blockade currents that are well separated from baseline noise (e.g., A3A5; see Fig. 2 top left), whereas low-contrast traces exhibit blockade levels only marginally below a noisy baseline (e.g., ssDNA and POC; see Fig. 2 top right and Fig. 2 bottom left), making events harder to distinguish. Moderate contrast traces reveal properties in middle (e.g., *λ*-DNA; see Fig. 2 bottom right).

**FIG. 2:**
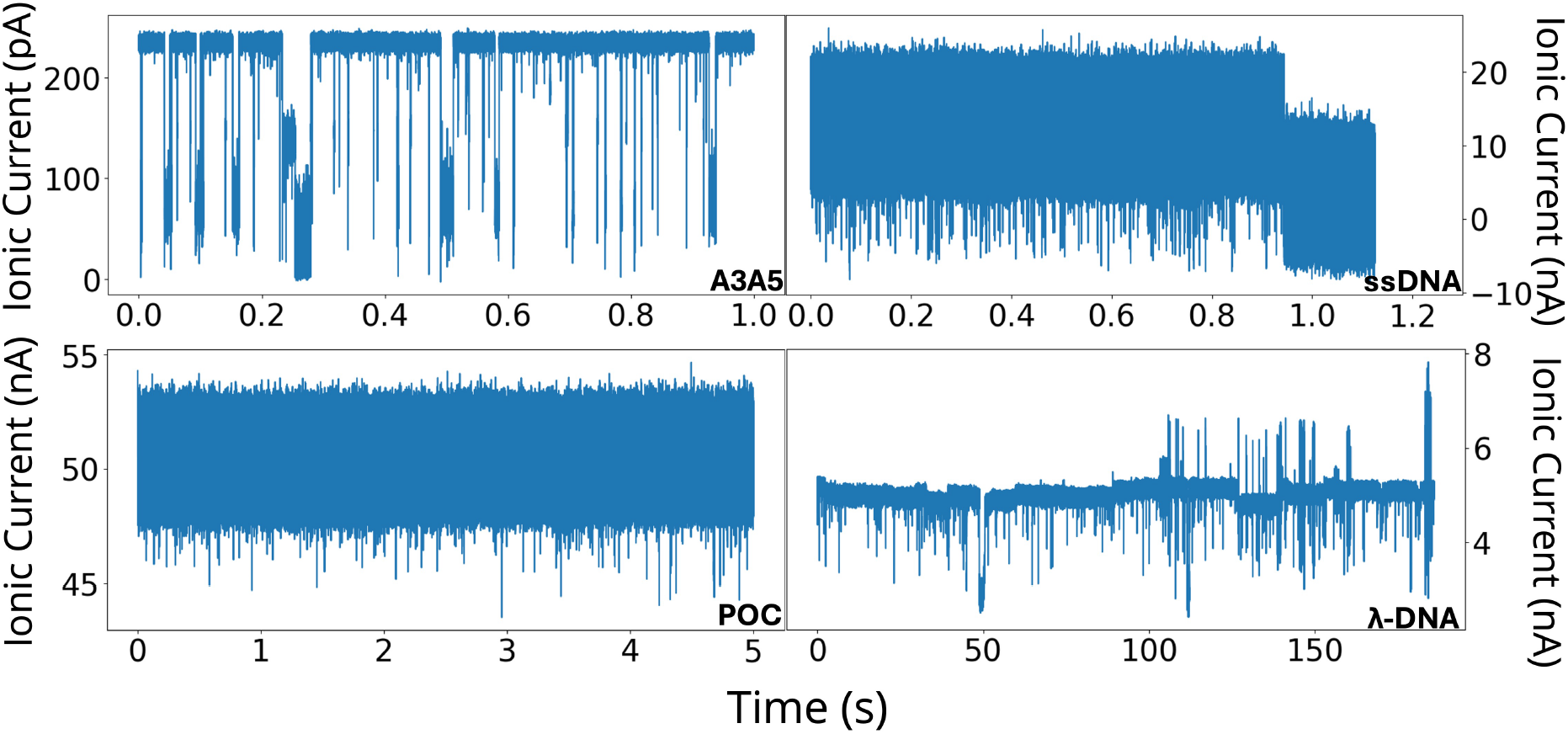
Time series of ionic current signals for the four distinct experiments in Table I: A3A5, ssDNA, POC, and *λ*-DNA, as denoted by the legends.

### D. Feature extraction

In order to read-out nanopore data, typically features need to be extracted in order to use these to train a model that later can predict the sequence of unknown analytes. It was shown that the height and mean in each current blockade include the essential information on the molecular identity and sequence [50]. Accordingly, after each event detection, we extract these features, as well as the dwell time, that is the duration of each event. An example of a current blockade and the corresponding extracted features is shown in Fig. 3.

**FIG. 3:**
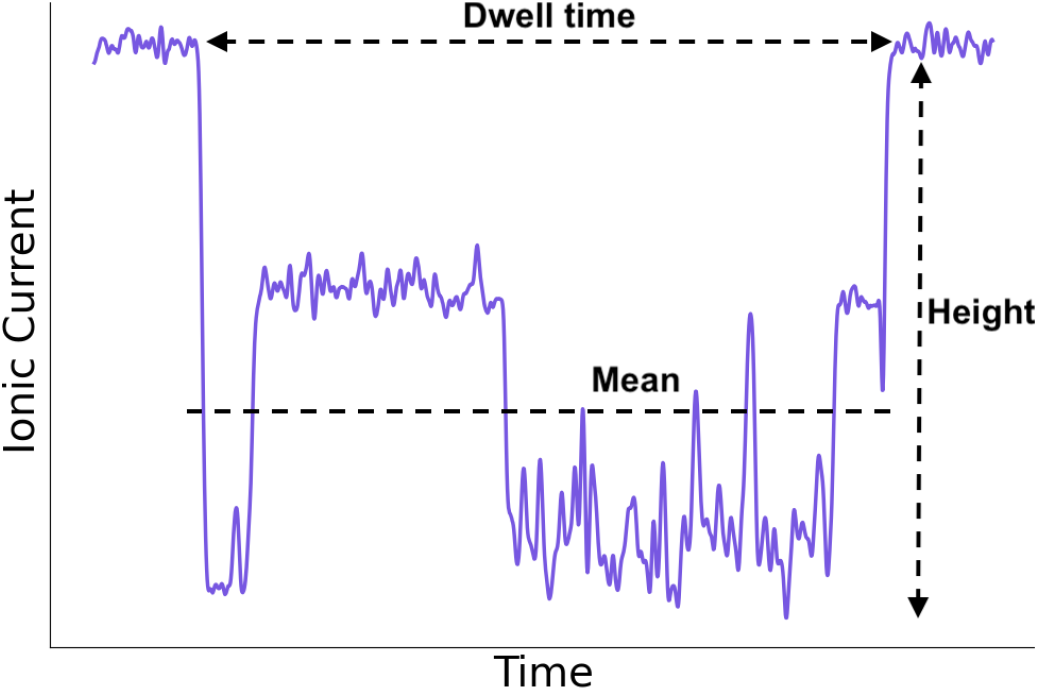
Example of an extracted event with labeled features. The event is baseline-corrected and the current blockage values are relative to the baseline. The mean value shows the average drop from baseline, not the average current. The three features that were extracted from all events are labeled and shown.

### E. Clustering algorithm

Once the features are extracted, these are clustered using the K-means [43] algorithms directly from the scikit-learn library [51] for further comparison. The K-means algorithm partitions data into a user-defined number of clusters by iteratively minimizing the within-cluster sum of squared distances from each point to its assigned cluster centroid. It assumes that clusters are spherical with uniform density, which may limit its effectiveness when clusters vary in shape or density [52].

**TABLE I:**
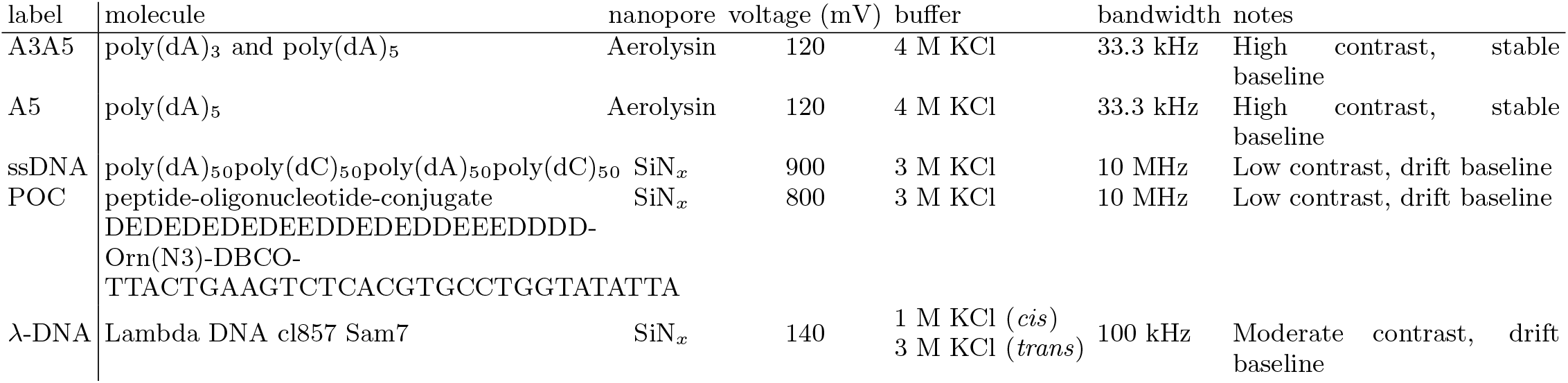
Overview of datasets analyzed in this work. “Label” is abbreviation for the experiment as used throughout; “molecule” describes the analyte; “nanopore” specifies pore type; “voltage” and “buffer” list the driving potential and electrolyte conditions; “notes” highlights signal-to-baseline contrast and baseline characteristics.

**TABLE II:** The number of events detected with each algorithmic scheme.

| algorithm | experiment |  |  |  |
| --- | --- | --- | --- | --- |
|  | A3A5 | ssDNA | POC | λ-DNA |
| CUSUM | 1030 | 149 | 1542 | 165 |
| PELT | 2354 | 156 | 1542 | 172 |
| CBED | 1963 | 159 | 1421 | 175 |

## III. RESULTS & DISCUSSION

### A. Event detection: comparison

We systematically evaluate and compare the performance of the three algorithms, CUSUM, PELT, and the enhanced cluster-based method (CBED) based on:

1. the numbers and characteristics of detected events,
2. the effectiveness of feature extraction and clustering,
3. the flexibility to adapt to varying experimental conditions, and (4) robustness to noise and signal variability. Using diverse experimental nanopore datasets, this study provides a detailed, side-by-side comparison that clarifies the practical strengths and limitations of each approach under different signal conditions.

We begin the analysis with the results on the event detection of the three schemes, CUSUM, PELT, and CBED, on all data mentioned above. Figure 4 shows the clustering of baseline versus blockade currents for each dataset: A3A5 exhibits clear separation of the two Gaussians, so the standard CBED thresholding is sufficient. Low signal-to-baseline ratio in ssDNA leads to overlapping clusters, low contrast combined with baseline drift in POC prevents clear separation, and baseline drift in lambda obscures moderate contrast, yielding indistinct clusters. Fixed-threshold detection is therefore ineffective, so we apply sliding-window adaptive threshold ex-tension in our enhanced CBED. Tab. II summarizes the number of events detected.

**FIG. 4:**
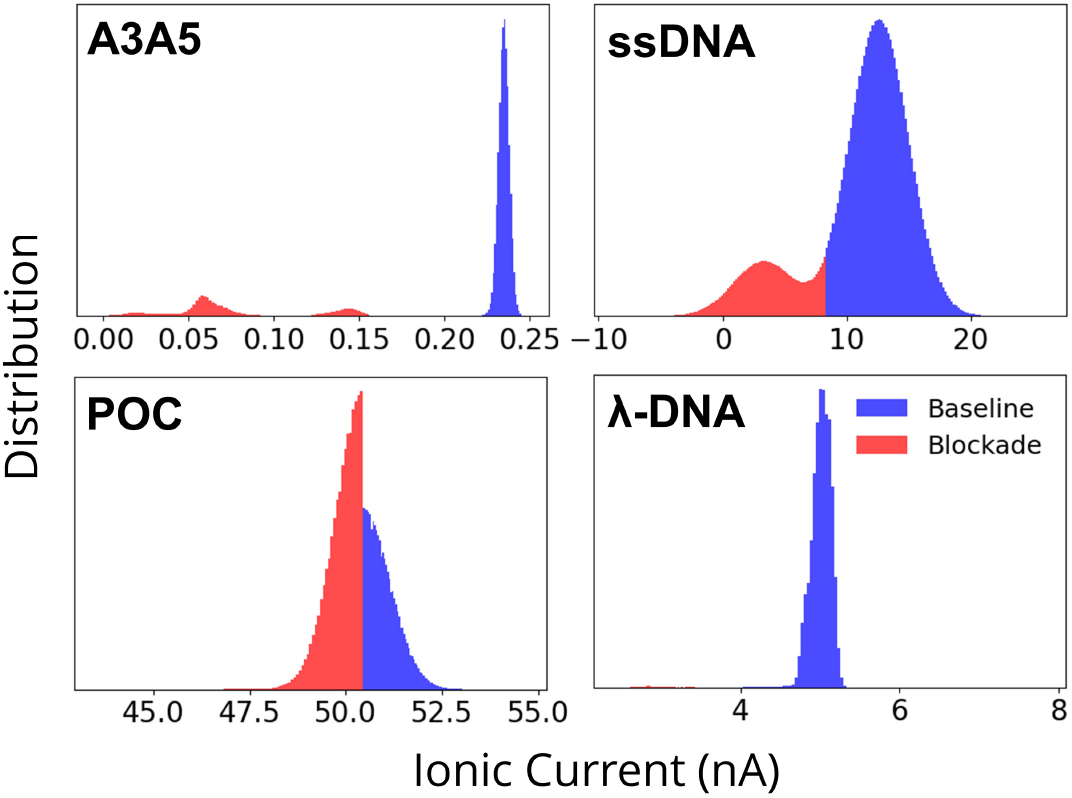
The cluster-based distributions of the ionic current signals for the four experiments (see legends) using the GMM algorithm. The distributions are assigned to blockade events (“Blockade”) and non-events (“Baseline”), respectively.

Across datasets, the relative performance of the three schemes is mainly determined by the dwell time variability and signal contrast. In the A3A5 dataset, where event durations span a wide range, CUSUM misses nearly half of the events detected by PELT, indicating reduced robustness when dwell times are heterogeneous. In contrast, CBED consistently captures both short and long events, reducing the number of events missed and enabling more reliable monitoring when event lengths are unpredictable. In the ssDNA dataset, all three methods produce similar event counts, with CBED detecting only slightly more events than PELT and CUSUM, suggesting that its increased sensitivity does not lead to substantial overcounting under low contrast. In the low contrast POC dataset, characterized by small blockade amplitudes and elevated baseline noise, CBED reports slightly fewer (8%) events than PELT or CUSUM. This decrease is consistent with improved rejection of noise induced detections rather than a loss of true blockades, although it also reflects the possibility that false positives become more likely under particularly challenging signal to noise conditions. For the moderately contrasted *λ*-DNA dataset, the experimental traces exhibit baseline drift, making a single fixed threshold unsuitable. We therefore incorporated adaptive thresholding into CBED to follow local baseline fluctuations as shown in Fig. 5. With this modification, CBED detects 175 events compared to 172 for PELT and 165 for CUSUM, with the modest increase attributable to occasional noise excursions crossing the adaptive threshold. We later show that these additional events form a separate cluster in the mean-height feature space and can be removed during downstream analysis. Overall, CBED matches or surpasses the sensitivity of established change-point methods, excelling in both high-contrast and difficult traces without sacrificing specificity in simpler cases.

**FIG. 5:**
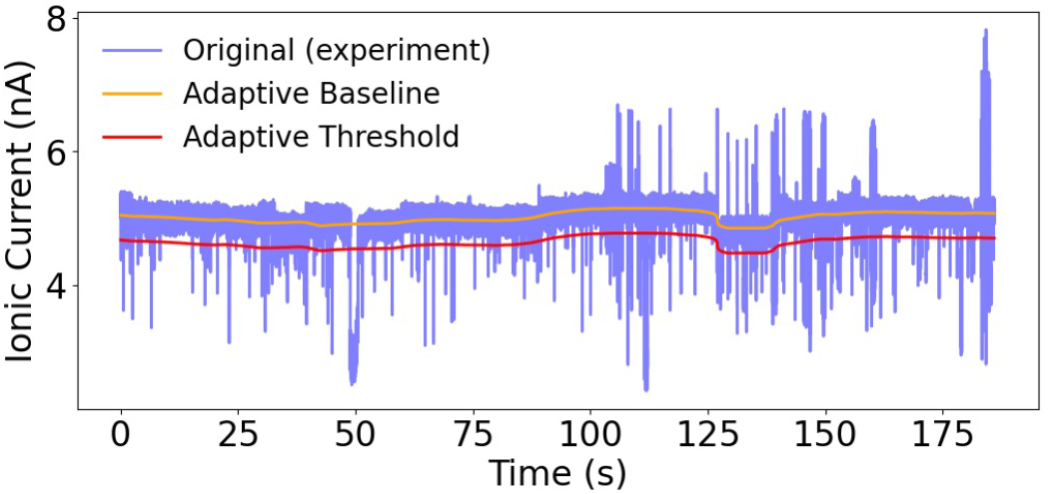
Ionic current trace for *λ*-DNA. The raw signal (blue) is overlaid with the adaptive baseline (orange) and its corresponding adaptive detection threshold (red). The baseline tracks slow drift in the open pore current, while the threshold follows the baseline to consistently detect downward blockade events by the CBED adaptive-threshold extension.

### B. Features and clustering

For all the events detected with the three algorithms and all four datasets, we extract the two features: mean blockade amplitude and blockade height and perform a clustering analysis. These two amplitude descriptors are widely used in nanopore signal analysis and have been shown to capture most of the discriminative information in blockade signals [53]. We next applied 2D K-means clustering for mean and height, which revealed clear groupings corresponding to baseline noise and real events of varying amplitudes. This perspective helps explain differences in event counts, because it shows whether additional detected events form coherent clusters or accumulate near the noise region. Across datasets, the CBED generally generates a tighter structure with less occupancy close to zero, while PELT and CUSUM more often retain shallow, near baseline modes that are consistent with noise.

This trend is most evident for the A3A5 dataset. In the height histogram (Fig. 6 left), four peaks are labelled i to iv at approximately 0.025, 0.1, 0.2, and 0.23 nA respectively. CBED resolves three peaks at positions ii, iii, and iv, with essentially no counts at lower heights. PELT and CUSUM reproduce these three peaks but show an additional shallow peak i, which is absent in CBED. In the mean histogram (Fig. 6 right), CBED shows two clean peaks at approximately −0.17 and −0.09 nA with no density near zero current, while PELT and CUSUM both display an extra peak close to zero at around −0.02 nA. To assign these peaks to the underlying analytes, we used an A5 only control dataset (Fig. 6 insets). The A5 control reproduces both peak iii and peak iv in the height histogram, with peak iv being the dominant mode, and shows a single peak at the deeper position in the mean histogram. Peaks iii and iv therefore both correspond to A5 blockades, with peak iv representing the main A5 population and peak iii likely reflecting an alternative A5 conformation, such as a partially folded or differently threaded state that produces a slightly shallower blockade. Peak ii, which is absent in the A5 control, is associated with A3. Peak i appears only in PELT and CUSUM and is most likely noise. The same logic applies to the mean histogram, where the peak around −0.17 nA corresponds to A5, the peak around −0.09 nA to A3, and the near zero peak seen only in PELT and CUSUM reflects false positive events.

**FIG. 6:**
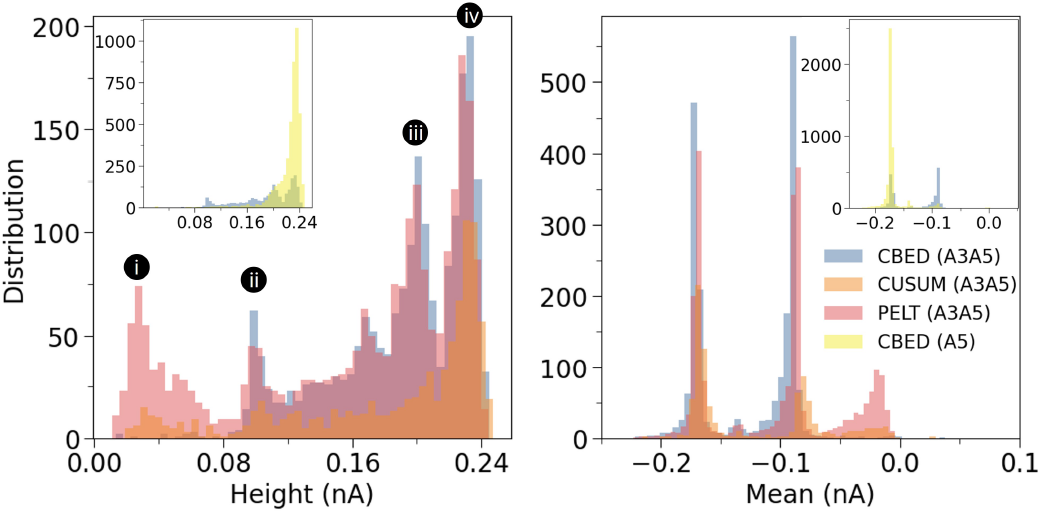
Distributions of extracted event features for A3A5 (main panel) and A5 (insets), obtained using the three detection schemes: CBED (blue), CUSUM (orange), and PELT (red). Left: blockage height. Right: mean blockade amplitude.

The same separation is reflected in the two dimensional clustering. In the mean height plane, CBED (Fig. 8 top left, blue) cleanly separates true A3 and A5 events into two tight groups, leaving almost no points in the near zero noise region, as expected for a mixture of A3 and A5 homopolymers. PELT (red) produces substantially overlapping clusters and a dispersed near zero group, consistent with undercounting of A3 blockades and numerous false positives. CUSUM does show the A3 related height feature near peak ii, but it is much weaker than in CBED and PELT, and the corresponding mean feature is also weak, indicating that fewer mid level A3 events are captured.

In ssDNA, the feature distributions and clusters align closely across all three methods, which supports the conclusion that CBED’s sensitivity does not lead to systematic over detection in low contrast data. The CBED height histogram (Fig. 7 left) closely matches PELT and CUSUM, with only a marginal shift toward the upper end of the 12 to 18 nA range, and the mean histograms (Fig. 7 right) overlap almost exactly. Consistent with this, the two dimensional clustering (Fig. 8 top right) shows CBED’s clusters directly on top of those from PELT and CUSUM, without an forming an additional noise cluster. At the other extreme, the low contrast POC dataset highlight how cluster compactness changes under strong baseline noise and very short dwell times. All methods detect a tight height distribution centered around 3 nA (Fig. 7 left), but CBED produces the sharpest peak and narrowest spread, while PELT and CUSUM show broader, more flat curves. The mean histograms (Fig. 7 right) show the same pattern, with CBED concentrated near −1.8 nA and PELT and CUSUM events spreading more symmetrically with longer tails towards zero blockade. In mean height space, CBED (Fig. 8 (bottom left), blue) forms a more compact cluster with less mixing into the noise region, consistent with improved rejection of noise induced event detection in this short event regime. For *λ*-DNA, the height spans roughly 0.5 to 2.5 nA for all algorithms, but PELT and CUSUM exhibit a small additional peak around 0.25 nA that is likely noise and is absent in the height histogram for the CBED events. The mean histograms peak between −0.5 and −0.8 nA, while CBED also shows a small extra peak at zero nA. Inspection of the traces indicates that this mode arises from rapid, large current swings that cross the threshold but cancel in the average (data not shown). In the mean height plane (Fig. 8 (bottom right), blue), these points form a distinct cluster near zero mean current despite substantial height variation. These can thus be removed during post processing if required.

**FIG. 7:**
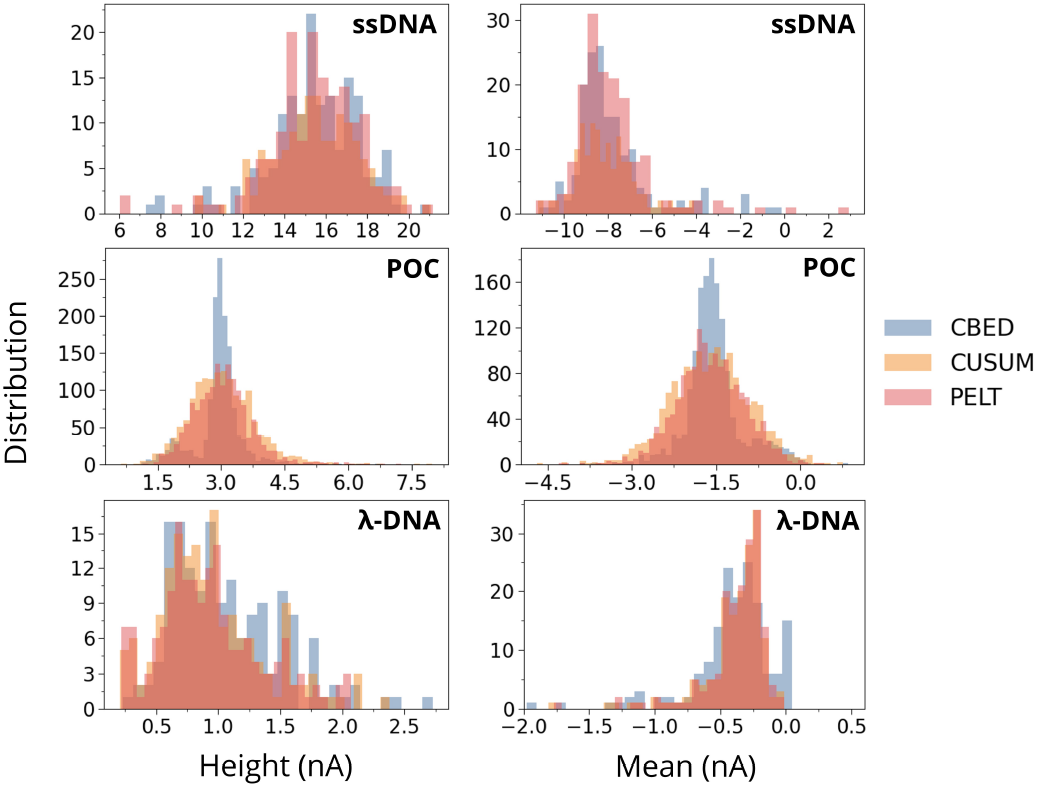
Distributions of extracted event features (left: blockage height, right: mean blockade amplitude) for the ssDNA, the POC, and the *λ*-DNA dataset, detected by CBED (blue), CUSUM (orange), and PELT (red), as denoted by the legends.

**FIG. 8:**
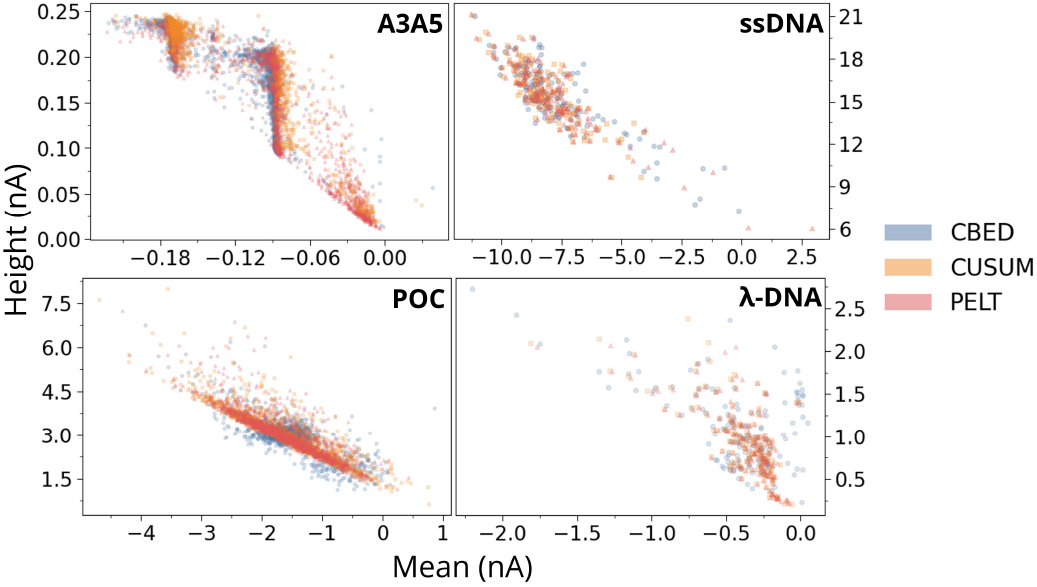
K-means clustering of the mean and height features of all events for the four datasets (A3A5, ssDNA, POC, and λ-DNA), as detected using all three event detection schemes(CBED (blue), CUSUM (orange), and PELT (red)).

### C. Flexibility and robustness

Finally, we compare the flexibility in terms of usage of the empirical parameter screening, as well as the computational performance of all three detection schemes. PELT requires the selection of a penalty term “pen” to balance detection sensitivity against over-segmentation. Choosing a proper “pen” typically involves testing several values and evaluating their impact on event counts. CUSUM is even more parameter-demanding: the implementation used here requires three tunable parameters: delta (the minimum change magnitude), hbook (the decision threshold), and sigma (the expected noise standard deviation). All three parameters must be set by trial and error to achieve a good trade-off between missed detections and false events. On the other hand, CBED requires only a simple vertical threshold between baseline and blockade events and a horizontal threshold for the duration. Specifically, the vertical threshold is determined directly from the clustering output by placing it at the valley between the two peaks, with no further calibration needed. If a drifting baseline makes a single global threshold unsuitable, the adaptive CBED extension recalculates the threshold locally, restoring accurate event calling automatically. This minimal need for parameter screening and its built-in ability to adjust automatically to baseline shifts make CBED significantly more user-friendly and robust than classical change-point detectors. In order to compare and support the handiness and efficiency of the event detection schemes, in Tab. III we report the computational times (in seconds) for the event detection. For the A3A5 dataset, we used the standard CBED implementation. In this case, CBED completes the detection in roughly 15 s; the fastest among the three methods (roughly 124 s for CUSUM and 20 s for PELT). For the ssDNA, POC, and *λ*-DNA datasets, the observed baseline shifts necessitate the use of the adaptive threshold extension of the CBED. This extension required roughly 177 s, 657 s, and 101 s, respectively, which exceeds the runtimes of CUSUM and PELT. Still its ability to adjust thresholds automatically in response to baseline shifts is an important advancement counter-balancing the additional computational time. Overall, these results demonstrate that standard CBED delivers superior speed on high-contrast data, while the adaptive extension provides the robustness needed for accurate event detection in the presence of baseline drifts, at the cost of increased runtime.

**TABLE III:** The computational time (s) of the three event detection schemes for all experiments considered here. The data for the CBED scheme with and without threshold adaptivity are shown for comparison.

| algorithm/experiment | A3A5 | ssDNA | POC | λ-DNA |
| --- | --- | --- | --- | --- |
| CUSUM | 123.9 | 12.5 | 166 | 32.3 |
| PELT | 20.5 | 2.9 | 119.7 | 22.8 |
| CBED (GMM) | 14.7 | - | - | - |
| CBED (GMM+adapt. thresh.) | - | 177.5 | 657.2 | 100.9 |

## IV. CONCLUSIONS

In order to improve the event detection in nanopore data, that is time series of an ionic current, in view of efficiency and speed, we have compared the performance of three algorithms. Unlike conventional change-point methods such as CUSUM and PELT, which rely on user-defined sensitivity parameters and sequential or penalty driven segmentation, CBED derives its detection threshold directly from the statistical structure of the signal itself. Specifically, a two-component Gaussian mixture model is fitted to the empirical current distribution, and the detection threshold is placed at the intersection of the two component densities corresponding to the natural valley separating open pore baseline from blockade events. This data-driven threshold determination eliminates the need for manual calibration and makes the algorithm inherently adaptive to the signal characteristics of each individual dataset. A strength of the algorithm is the on-the-fly adaptivity for baseline drifts.

The comparison of the three event detection algorithms reveals that CBED performs consistently across datasets with very different signal type. For high-contrast traces, it detects events reliably and with low computational cost. It also requires less parameter tuning than change-point based approaches. For moderate-contrast signals, it remains competitive and gives stable event statistics. For low-contrast or drifting baselines, mainly in solid-state nanopores, the adaptive version improves detection because it avoids using a single global threshold. In addition, the detected events produce cleaner feature distributions, with fewer near-baseline false detections that can otherwise blur clustering results. Overall, the results show that the proposed framework is a practical alternative to standard schemes. It balances accuracy with speed, and it supports downstream analysis by providing more consistent events.

Each of the three methods has its own strengths. CUSUM remains the natural choice when real-time or streaming analysis is required, though its performance is the most sensitive to parameter selection, since all three of its thresholds must be set by the user typically based on trial and error. PELT offers a robust and largely parameter-free alternative for offline work, with only a single penalty term to adjust, and it is particularly efficient on low-contrast or slowly drifting data where its batch segmentation strategy is well-suited. CBED sidesteps manual threshold tuning entirely and handles baseline drift through its adaptive fitting, at the cost of somewhat higher computation on challenging datasets. Taken together, the event detection algorithmic comparison emphasizes that none of the algorithms is universally superior: the most appropriate choice depends on the characteristics of the recorded signals and whether the application demands online processing or can afford the greater flexibility of an offline pipeline.

## ACKNOWLEDGMENTS

PW, MK, and MF are thankful to Angel Diaz-Carral, Martin Rotegui, Ayberk Koc, and Ana Poput for useful discussions. The computing time provided in the NHR Center NHR4CES at RWTH Aachen University (project number p0022407 and p0021740) is greatly acknowledged. This is funded by the Federal Ministry of Education and Research, and the state governments participating on the basis of the resolutions of the GWK for national high performance computing at universities.

This work is part the nanodiagBW consortium (project number 03ZU1208BI, 03ZU1208AJ and 03ZU1208BG) funded by the Federal Ministry for Research, Technology and Space (BMFTR) within the Clusters4Future initiative. Funding from the German Funding Agency (Deutsche Forschungsgemeinschaft) and DFG - Project number 508324943 is greatly acknowledged. C-YL, KK, and MD greatly acknowledge the National Institutes of Health (NIH) for funding through the grant NIH R01HG012413. MM and PDJ acknowledge financial support from the State Ministry of Baden-Wuerttemberg for Economic Affairs, Labour and Tourism.

## V. DATA AVAILABILITY STATEMENT

The data used for the analysis can be shared upon reasonable request. The Machine Learning workflow in this work is available on GitHub.

Experimental data for A3A5 were recorded by T.E. in the laboratory of J.C.B. under a former affiliation.

