## Supplemental Figures for "Nanopore event detection in a simple and adaptive way"

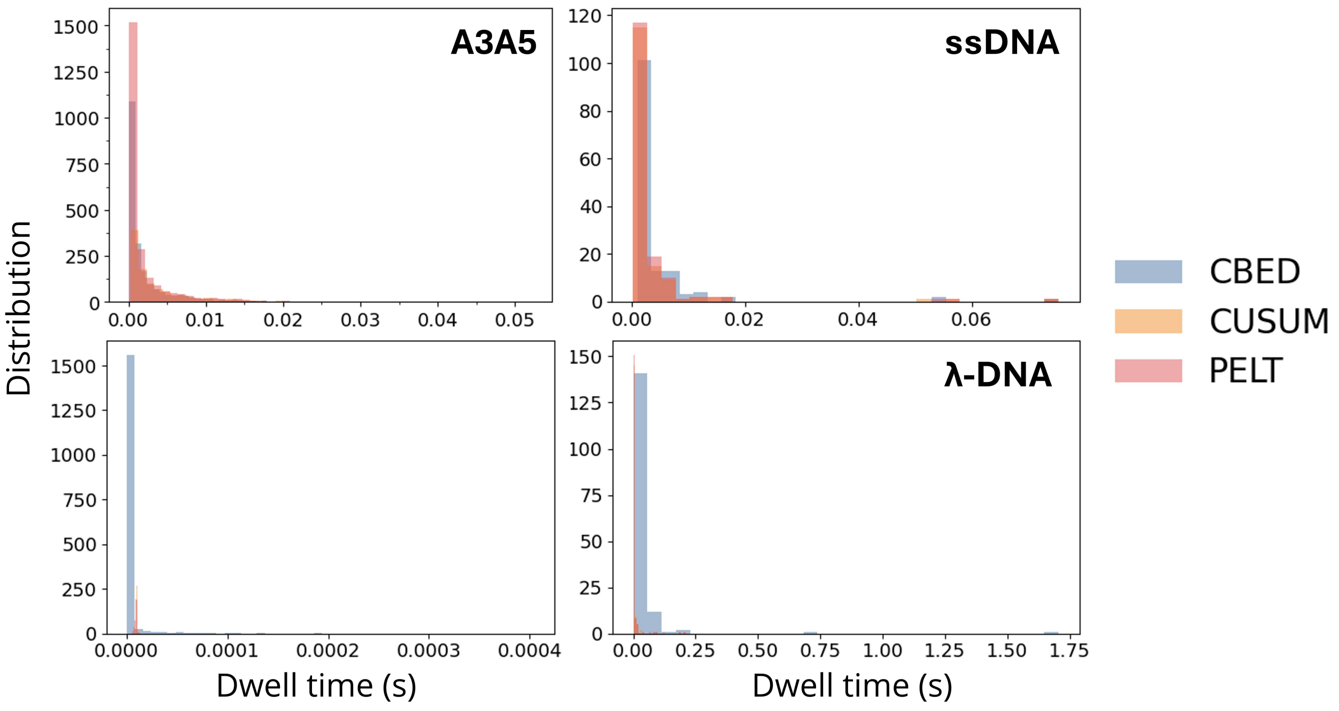


FIG. 1. Distributions of extracted event features dwell time for the A3A5, the ssDNA, the POC and the λ-DNA dataset, detected by CBED (blue), CUSUM (orange), and PELT (red).

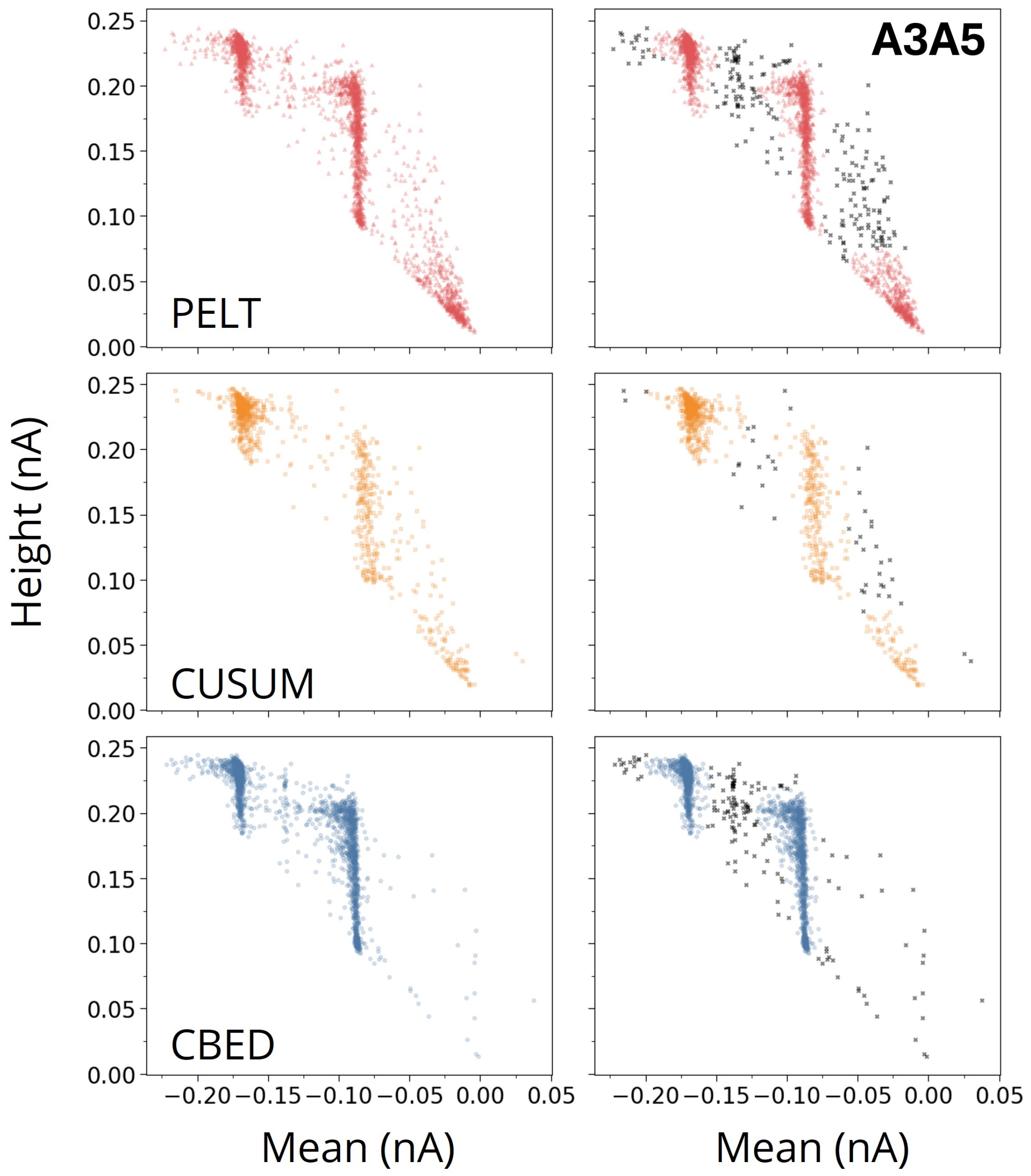


FIG. 2. Mean–depth clustering of A3A5 events. Results from K-means (left) and DBSCAN (right) for events detected by PELT, CUSUM, and CBED.

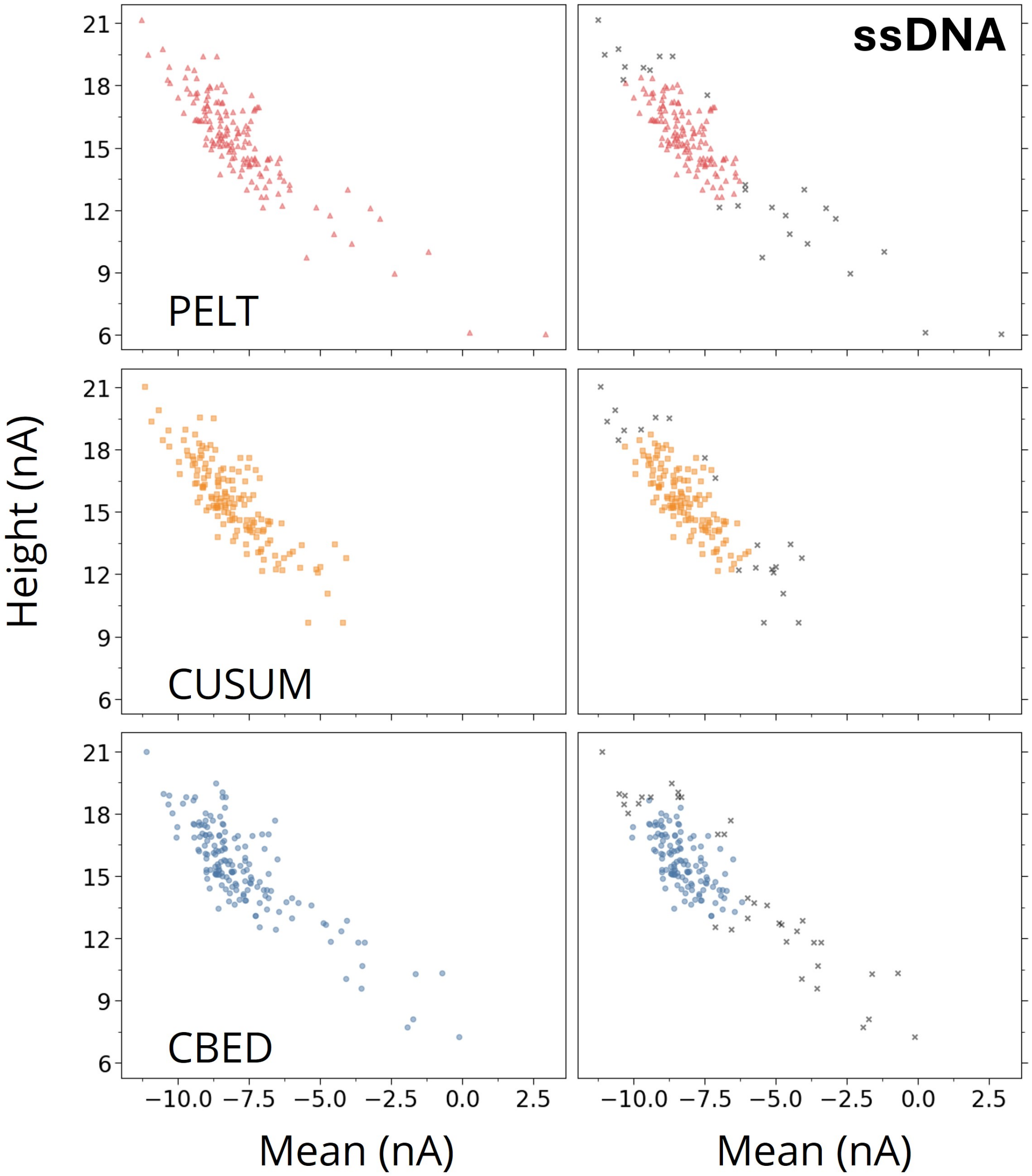


FIG. 3. Mean-depth clustering of ssDNA events. Results from K-means (left) and DBSCAN (right) for events detected by PELT, CUSUM, and CBED.

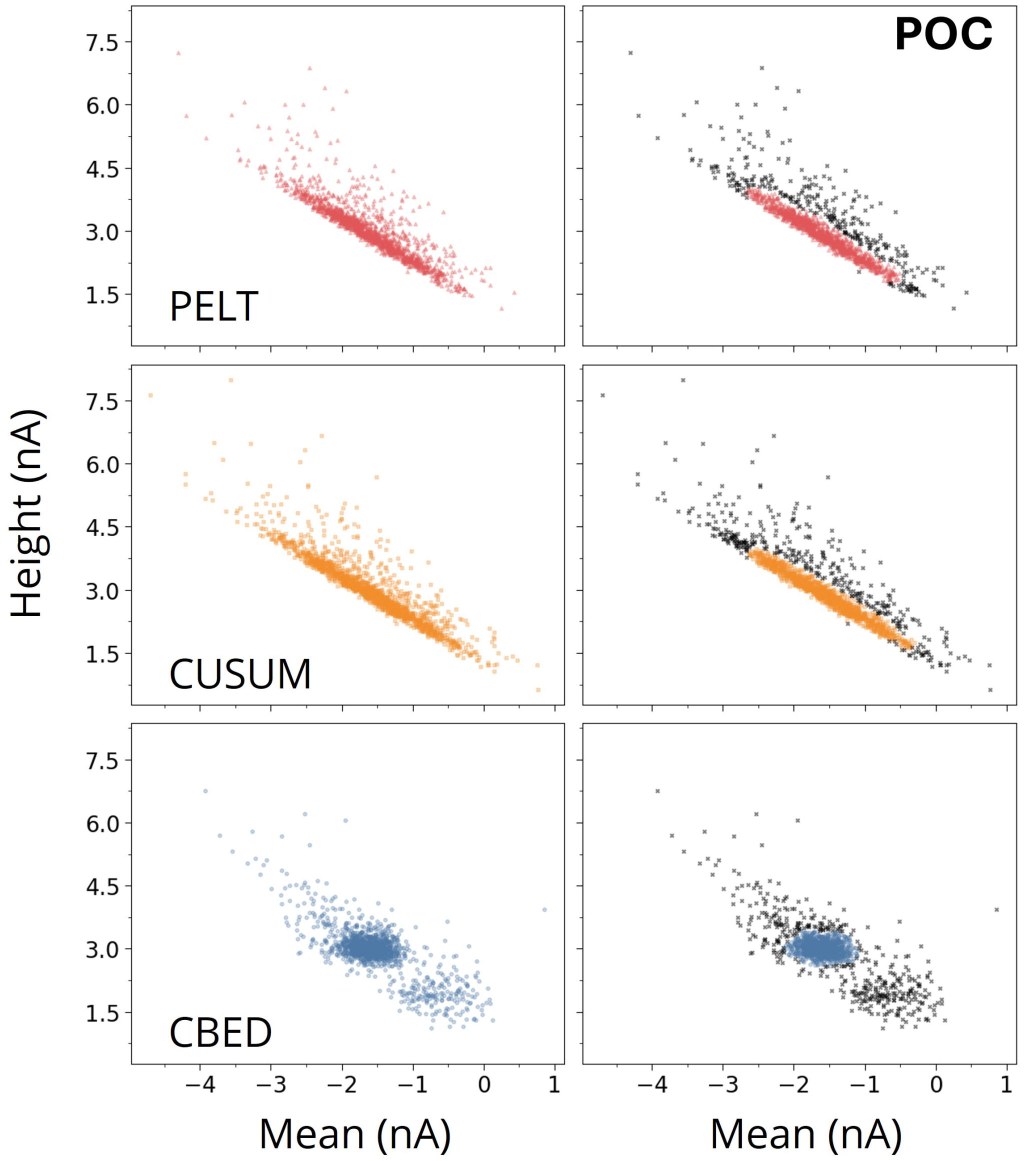


FIG. 4. Mean-depth clustering of POC events. Results from K-means (left) and DBSCAN (right) for events detected by PELT, CUSUM, and CBED.

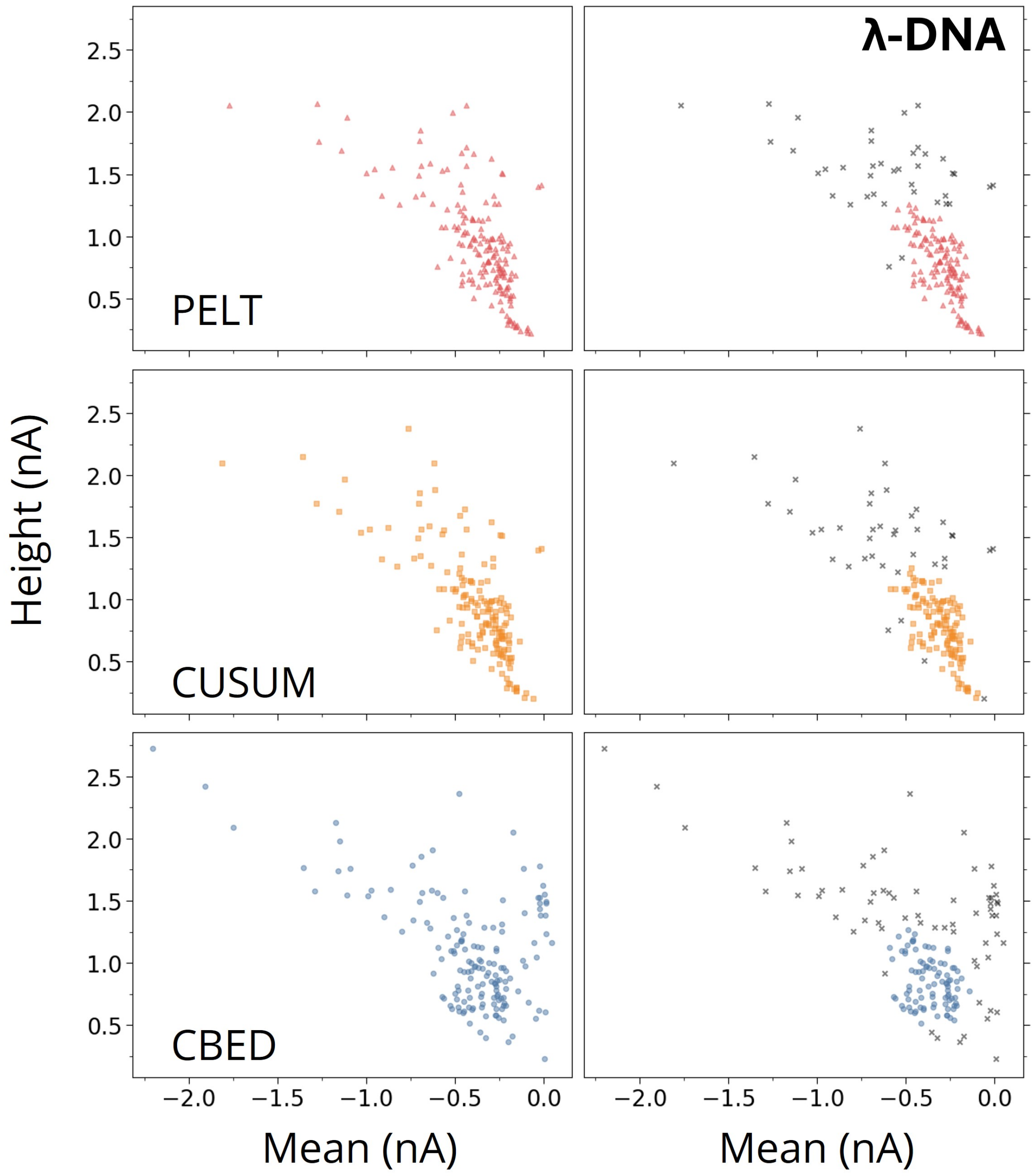


FIG. 5. Mean-depth clustering of $\lambda$-DNA events. Results from K-means (left) and DBSCAN (right) for events detected by PELT, CUSUM, and CBED.

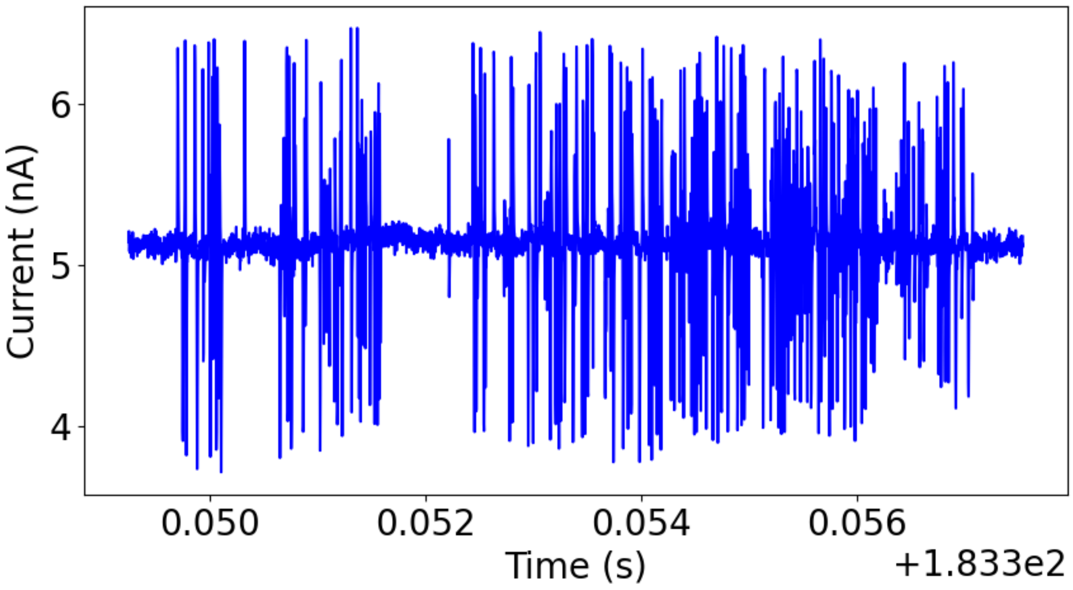


FIG. 6. Noise_$\lambda$-DNA_CBED.
